# Sterilizing Effects of Novel Regimens Containing TB47, Clofazimine and Linezolid in a Murine Model of Tuberculosis

**DOI:** 10.1101/2021.04.04.438375

**Authors:** Wei Yu, Yusuf Buhari, Shuai Wang, Xirong Tian, H. M. Adnan Hameed, Zhili Lu, Gift Chiwala, Md Shah Alam, Gregory M. Cook, Dmitry A. Maslov, Nanshan Zhong, Tianyu Zhang

## Abstract

Toxicity and inconvenience associated with the use of injectable drug-containing regimens for tuberculosis (TB) have made all-oral regimens a preferred alternative. Widespread resistance to fluoroquinolones and pyrazinamide makes it essential to identify new drug candidates and study their effects on current regimens for TB. TB47 is a pyrazolo[1,5-a]pyridine-3-carboxamide with powerful synergistic *in vitro* and *in vivo* activities against mycobacteria, especially with clofazimine. Here, we investigated the bactericidal and sterilizing activities of novel oral regimens containing TB47 + clofazimine + linezolid, and the potential roles of levofloxacin and/or pyrazinamide in such drug combinations. Using a well-established mouse model, we assessed the effect of these regimens on bacterial burden in the lung during treatment and relapse (4 months after stopping treatment + immunosuppression). Our findings indicate that the TB47 + clofazimine + linezolid + pyrazinamide, with/without levofloxacin, regimens had fast-acting (4 months) sterilizing activity and no relapse was observed. When pyrazinamide was excluded from the regimen, treatment times were longer (5-6 months) to achieve sterilizing conditions. We propose that TB47 + clofazimine + linezolid can form a highly sterilizing block for use as an alternative pan-TB regimen.

## INTRODUCTION

Although tuberculosis (TB) is a treatable infectious disease, in 2019, an estimated 10 million people developed active TB, with 1.2 million deaths as well as 208,000 deaths among HIV/TB co-infected individuals (1). For patients with drug-susceptible TB, standard treatment with the four powerful first-line anti-TB drugs results in acceptable cure rates. Patients who are infected with strains resistant to at least rifampicin (RIF) and isoniazid, called multidrug-resistant TB (MDR-TB), possess limited treatment alternatives and are practically incurable by the standard first-line treatment. Approximately 500,000 new cases of MDR-TB occur annually with a low treatment success rate at ≤ 57% (1). It usually takes 9 - 12 months for regimens containing at least 4 - 6 drugs, including at least one injectable agent, to treat patients with MDR-TB (2). More seriously for patients, extensively drug-resistant TB (XDR-TB) is MDR-TB plus resistance to a fluoroquinolone (FQ) and an injectable agent, with an unacceptably low treatment success rate of only ≤ 39% (3). Novel drugs and regimens to shorten and improve MDR/XDR-TB treatment success are urgently needed. If such regimens do not contain RIF and isoniazid, they may be applicable to both drug-susceptible and MDR-TB. Furthermore, if injectable drugs and FQs are also not included in the combination, such regimens may be useful for XDR-TB treatment. For instance, the recent FDA-approved combination, BPaL (bedaquiline + pretomanid + linezolid), is a fully oral regimen with relatively shorter course of treatment that has shown remarkably promising clinical effect against both MDR- and XDR-TB with manageable side effects. More of such regimens are desired to improve TB treatment.

TB47, a pyrazolo[1,5-a]pyridine-3-carboxamide, is a promising anti-mycobacterial agent as its mechanism of action is not shared with other existing drugs (4). It can block the cytochrome *bc*_*1*_-*aa*_*3*_ oxidase complex by targeting the QcrB subunit, thereby causing a reduction in intracellular ATP and eventually inhibiting the growth of mycobacteria (5,6). Although mono-therapy with TB47 is rather bacteriostatic than bactericidal in mouse TB models due to rerouting of energy metabolism from QcrB to the cytochrome *bd* oxidase (7,8), it has been proved to provide good synergistic effects in combination with either pyrazinamide (PZA) or RIF, demonstrating a potential role in combination therapies (6). With cytochrome *bd* inactivation, TB47 holds a significant bactericidal potential (6). We revealed that TB47 exerted high bactericidal and sterilizing activity and corresponding remarkable treatment-shortening effect in mice infected with *Mycobacterium ulcerans* which lacks cytochrome *bd* activity naturally (5,7). It has also shown a very good synergistic activity with clofazimine (CLF) against *Mycobacterium abscessus* and *M. tuberculosis* both *in vitro* and *in vivo* (10,11), and even played a remarkable role in shortening MDR-TB treatment duration from ≥ 9 to 5 months in a well-established murine model of TB (11). CLF competes with menaquinone and kills bacteria by generating lethal levels of reactive oxygen species (12), and regimens containing CLF and cytochrome *bc*_1_ inhibitors or other electron transport chain-targeting drugs provide enhanced killing of both replicating and non-replicating *M. tuberculosis* (13–15), suggesting that dual or multiple targeting of the electron transport chain may be a novel approach to enhanced killing of mycobacteria (7,10).

Considering the extensive resistance to FQs (16) and PZA (17–19) by DR-TB, the serious side effects and inconvenience associated with injectable drugs, as well as the fact that the patent-expiring linezolid (LZD) showed a very good activity against DR-TB in clinical use (20), we evaluated the effect of partially or totally replacing them with LZD in drug combinations containing TB47 + CLF as a block to study their benefits in new regimens and to explore the possibility of developing a novel, fully oral pan-TB regimen.

## RESULTS

### Establishment of *M. tuberculosis* infection in female BALB/c mice

The scheme of the experimental design is shown in Table 1. On the day after infection (Day −15), *M. tuberculosis* H37Rv reached a cell density of 4.64 ± 0.19 mean log_10_ CFU (colony-forming unit) in the lungs. At the time of treatment initiation (Day 0), the *M. tuberculosis* burden increased to 8.18 ± 0.14 mean log_10_ CFU, demonstrating that a heavy infection was established in the lungs. All nine untreated mice (Table 1) succumbed to TB infection between day 22 and day 25 post-infection.

**Table 1.**
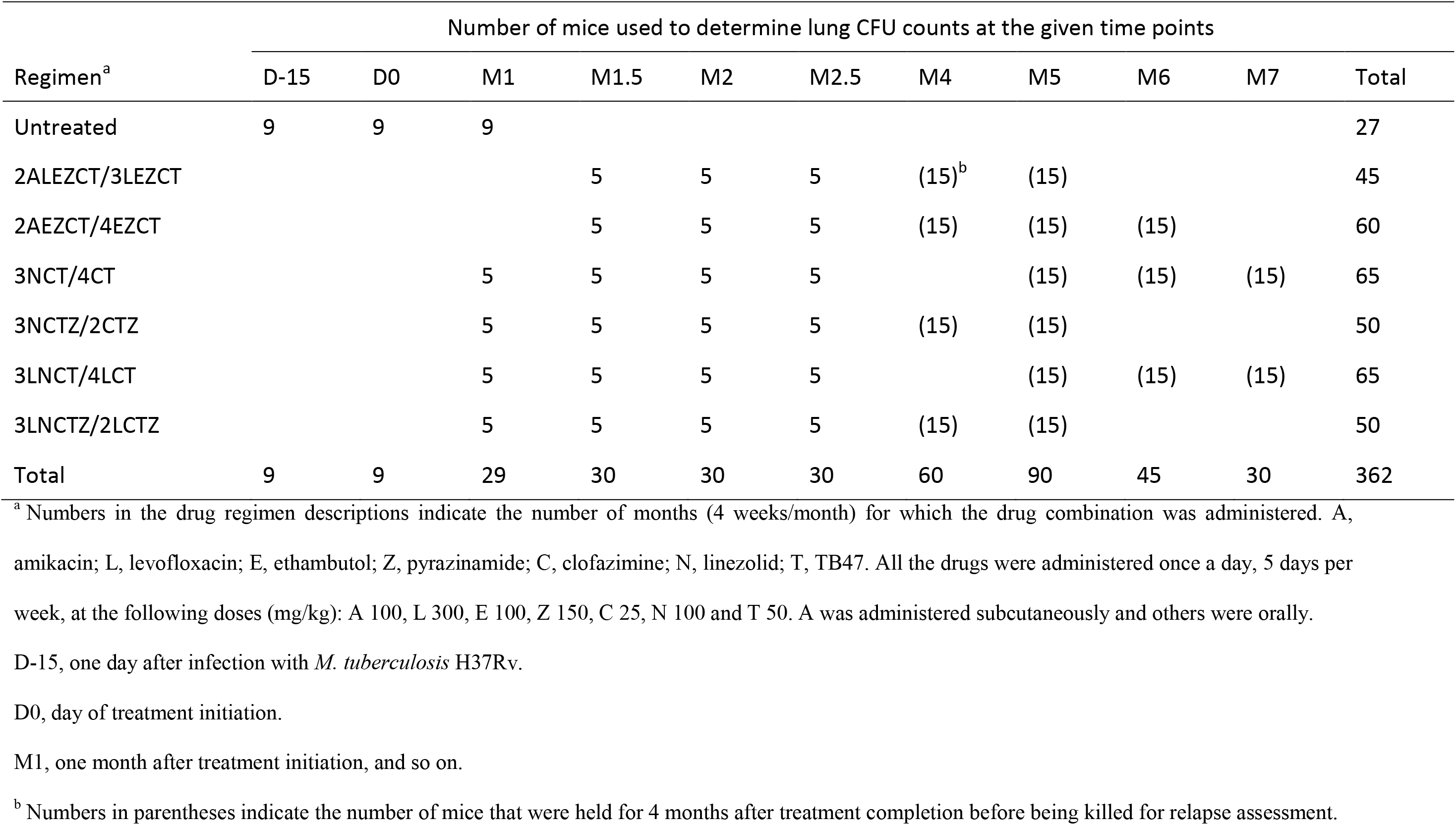
Scheme of the experiment.

### Bactericidal activities of the test regimens

Details of lung CFU counts obtained at various time points are presented in Table 2. While two of five mice in the NCTZ group, and only one of five mice in the LNCTZ group were culture negative (Table 2), all mice treated with other regimens remained culture positive after 2.5 months of treatment. The mean log_10_ CFU counts in the lungs declined significantly in all groups during the course of treatment (Table 2, Figure 1). LZD was used only for 3 months due to side effects (11). Addition of levofloxacin (LVX) to the AEZCT regimen resulted in stronger bactericidal activity compared to the AEZCT regimen alone (*P* = 0.0003 < 0.001) (Figure 1a). In contrast, contribution of LVX to the bactericidal activities of NCT and NCTZ was not significant (*P* > 0.05) (Figure 1b). However, there were significant differences between PZA-containing (NCTZ and LNCTZ) and corresponding PZA-lacking regimens (NCT and LNCT, *P* < 0.0001 and *P* < 0.001, respectively) (Figure 1b). Compared to ALEZCT, the stronger regimen containing amikacin (AMK), the injectable drug, NCTZ showed better bactericidal activity (*P* = 0.04 < 0.05), NCT showed less bactericidal activity (*P* = 0.02 < 0.05), LNCT and LNCTZ showed no difference (*P* = 0.097, *P* = 0.65, respectively) (Table 2).

**Table 2.**
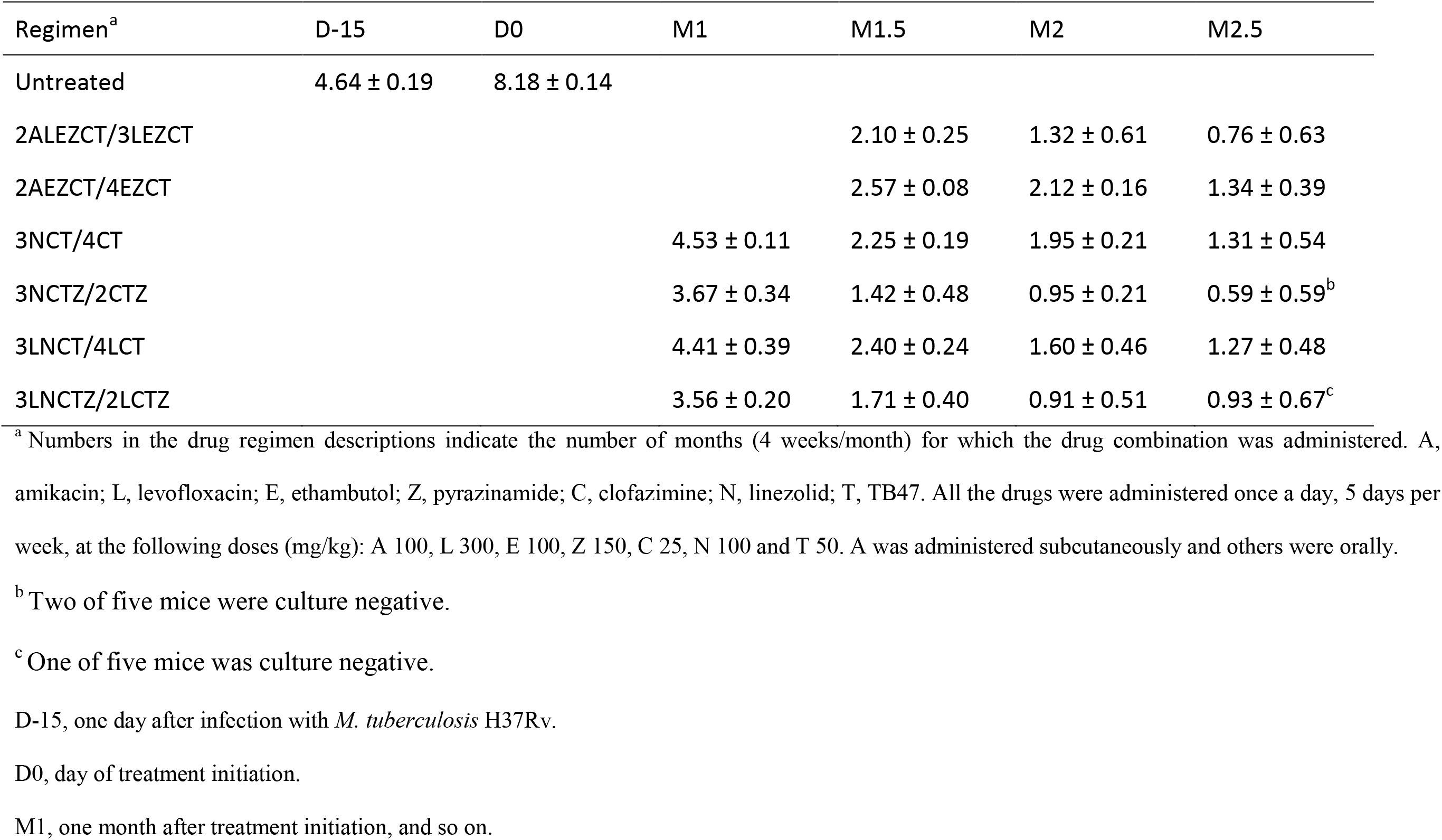
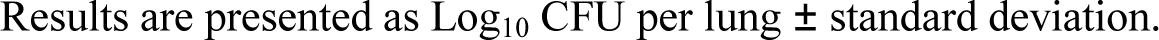
Lung CFU counts during the course of experiment.

| Regimen<sup>a</sup> | D-15 | D0 | M1 | M1.5 | M2 | M2.5 |
| --- | --- | --- | --- | --- | --- | --- |
| Untreated | 4.64 ± 0.19 | 8.18 ± 0.14 |  |  |  |  |
| 2ALEZCT/3LEZCT |  |  |  | 2.10 ± 0.25 | 1.32 ± 0.61 | 0.76 ± 0.63 |
| 2AEZCT/4EZCT |  |  |  | 2.57 ± 0.08 | 2.12 ± 0.16 | 1.34 ± 0.39 |
| 3NCT/4CT |  |  | 4.53 ± 0.11 | 2.25 ± 0.19 | 1.95 ± 0.21 | 1.31 ± 0.54 |
| 3NCTZ/2CTZ |  |  | 3.67 ± 0.34 | 1.42 ± 0.48 | 0.95 ± 0.21 | 0.59 ± 0.59<sup>b</sup> |
| 3LNCT/4LCT |  |  | 4.41 ± 0.39 | 2.40 ± 0.24 | 1.60 ± 0.46 | 1.27 ± 0.48 |
| 3LNCTZ/2LCTZ |  |  | 3.56 ± 0.20 | 1.71 ± 0.40 | 0.91 ± 0.51 | 0.93 ± 0.67<sup>c</sup> |

<sup>a</sup> Numbers in the drug regimen descriptions indicate the number of months (4 weeks/month) for which the drug combination was administered. A, amikacin; L, levofloxacin; E, ethambutol; Z, pyrazinamide; C, clofazimine; N, linezolid; T, TB47. All the drugs were administered once a day, 5 days per week, at the following doses (mg/kg): A 100, L 300, E 100, Z 150, C 25, N 100 and T 50. A was administered subcutaneously and others were orally.
<sup>b</sup> Two of five mice were culture negative.
<sup>c</sup> One of five mice was culture negative.
D-15, one day after infection with *M. tuberculosis* H37Rv.
D0, day of treatment initiation.
M1, one month after treatment initiation, and so on.
Results are presented as Log<sub>10</sub> CFU per lung ± standard deviation.

**Figure 1.**
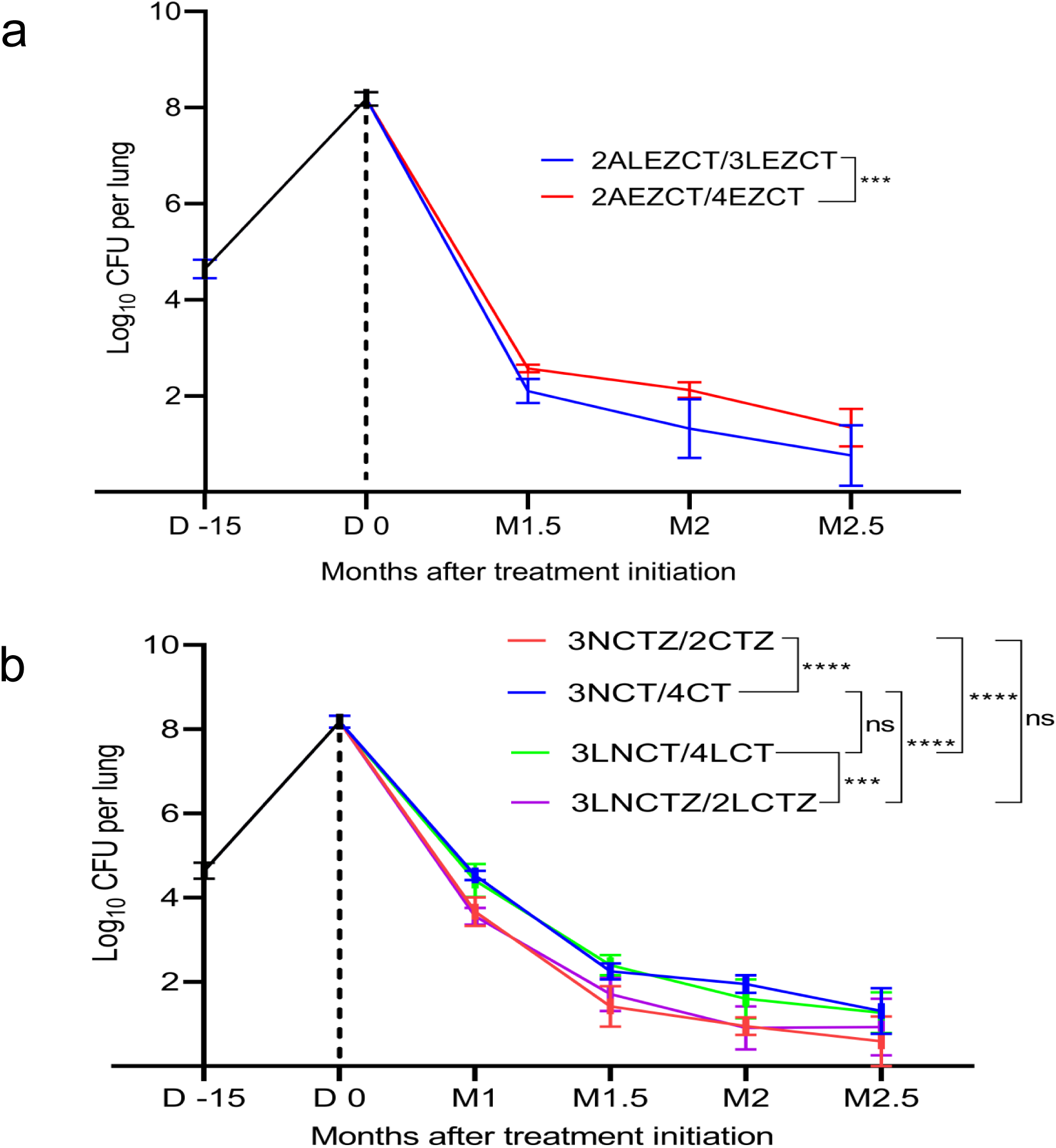
Bactericidal activities of the studied regimens as observed during the course of experiment. (a) Adding LVX to AEZCT had a noticeable effect on its bactericidal activity. (b) Addition of LVX did not result in significant contribution to activities of any of the NCT-based regimens. However, addition of PZA resulted in significant differences between PZA-containing and PZA-lacking regimens. Incorporation of both LVX and PZA enhanced the activity of NCT, but the activity of NCT with PZA alone equals or even surpasses the activity of NCT with both LVX and PZA added. ns, not significant, *P* > 0.05; ***, *P* < 0.001; ****, *P* < 0.0001. D −15, Day −15; M1, month 1, and so on.

### Relapse rates after treatment cessation

*M. tuberculosis*-infected mice were subjected to various duration of treatment using various combination regimens, after which a number of mice were held for 3.5 months followed by two weekly immunosuppression treatment with dexamethasone (DXM) prior to euthanasia and detection of *M. tuberculosis* in lungs for relapse (Table 1). Treatment with 2ALEZCT/2LEZC for 4 months resulted in a 13.33 % (2/15) relapse rate, which later reduced to 0 % (0/15) with another month of LEZC (Table 3). These results are almost the same with those from our previous study (11). In contrast, treatment with 2AEZCT/3EZCT for 5 months and 2AEZCT/4EZCT for 6 months yielded a 33.33 % (5/15) and 0 % (0/15) relapse rates, respectively (Table 3), which indicated that LVX played a role in shortening the treatment duration in ALEZCT combination, though it did not show a sharp activity in lowering the *M. tuberculosis* burden in lungs during the 2.5 months of treatment as mentioned above. Similarly, 3LNCT/2LCT for 5 months achieved 0% relapse rate, a month earlier than 3NCT/3CT did, which also suggested that LVX has played a role in the LNCT combination. Unexpectedly, after 4 and 5 months of treatment, both 3NCTZ/2CTZ- and 3LNCTZ/2LCTZ-treated mice showed 0 % (0/15) relapse rates (Table 3), which could not provide evidence of benefit of LVX in LNCTZ combination. This was consistent with the lack of difference in their bactericidal activities. We speculate that the actual cure time could be even shorter than 4 months for LNCTZ and NCTZ groups. Adding PZA into NCT significantly (*P* < 0.0001) shortened the treatment duration as 3NCTZ/2CTZ achieved 0 % (0/15) relapse rate at least 2 months earlier than 3NCT/4CT, which was accordant to the bactericidal activities of these two regimens. We could not provide evidence of benefit of PZA in the combination 3LNCTZ/2LCTZ as we had no cohort of mice relapsed in either 3LNCTZ/2LCTZ or 3LNCT/4LCT at designed time points. All four oral regimens could achieve 0 % (0/15) relapse rate in 4 to 6 months, which were shorter than or equal to the treatment duration of AMK-containing regimens (5 to 6 months). This indicated that LZD may be able to replace ALE or ALEZ in the control regimen 2ALEZCT/3LEZC and still provide equal or even better outcome.

**Table 3.** Proportion of mice relapsing after treatment completion.

| Duration of treatment<br>(months) | Regimen<sup>a</sup> | No. of culture-positive<br>mice/total No. of mice (% of<br>culture-positive)<sup>b</sup> |
| --- | --- | --- |
| 4 | 2ALEZCT/3LEZCT | 2/15 (13.33%)<br>[4, 51]<sup>c</sup> |
|  | 2AEZCT/4EZCT | 5/15 (33.33%)<br>[76, 88, 180, 550, 410]<sup>c</sup> |
|  | 3NCTZ/2CTZ | 0/15 (0%) |
|  | 3LNCTZ/2LCTZ | 0/15 (0%) |
| 5 | 2ALEZCT/3LEZCT | 0/15 (0%) |
|  | 2AEZCT/4EZCT | 5/15 (33.33%)<br>[291, 161, +++, +++, +++]<sup>c</sup> |
|  | 3NCT/4CT | 1/15 (6.67%)<br>[46]<sup>c</sup> |
|  | 3NCTZ/2CTZ | 0/15 (0%) |
|  | 3LNCT/4LCT | 0/15 (0%) |
|  | 3LNCTZ/2LCTZ | 0/15 (0%) |
| 6 | 2AEZCT/4EZCT | 0/15 (0%) |
|  | 3NCT/4CT | 0/15 (0%) |
|  | 3LNCT/4LCT | 0/15 (0%) |
| 7 | 3NCT/4CT | 0/15 (0%) |
|  | 3LNCT/4LCT | 0/15 (0%) |

<sup>a</sup> Numbers in the treatment regimen descriptions represent the number of months for which the drug combination was administered. Relapse was defined as ≥ one *M. tuberculosis* colony appeared on the plates spread with the entire lung homogenate 4 months after the indicated duration of treatment. Each lung homogenate was equally divided and plated on 4 plates.
<sup>b</sup> The proportion of mice with culture-positive relapse was determined by holding cohorts of mice for an additional 4 months after treatment completion.
<sup>c</sup> Values in brackets indicate the number of colonies from lung homogenates of mice with relapse after treatment completion.
+++ indicates too many to count.

## DISCUSSION

Potency of the standard short-course regimen consisting of first-line drugs (RIF + isoniazid + PZA + ethambutol (EMB)) for drug-susceptible TB requires at least 6 months of treatment. However, for DR-TB, the duration of treatment is even longer and treatment options are limited. Emergence of DR-TB, therefore, poses a significant threat to individual patient management and global TB control efforts. For this reason, a plethora of safer and accessible novel regimens that could shorten the duration of DR-TB treatment are urgently needed (21), one of which is the recent FDA-approved BPaL.

Injectable drugs are not recommended for TB treatment due to their toxicity, poor efficacy and causation of irreversible adverse effects, and may regularly need to be taken at healthcare facilities. The WHO suggests the phasing out of injectable drug-containing regimens and adoption of a shorter all-oral bedaquiline-containing regimen for treatment of MDR/RR-TB (22). All-oral regimens, therefore, as opposed to injectable drug-containing regimens, are a preferred alternative for TB treatment as shown by improved clinical outcomes. It has become quite important to study the synergistic activities of novel drugs/compounds with no cross-resistance with other anti-TB drugs as activities of some compounds may be determined by efficacies of their companion drug(s) (23).

TB47 is an antimycobacterial candidate that has a good safety profile and ideal pharmacokinetic parameters (5,6), as well as good synergistic effects with other TB drugs (6,10). We used the Bangladesh regimen + TB47 (ALEZCT) from our previous study (11) as a positive control here, in which we further confirmed the highly synergistic activity of TB47 + CLF in shortening MDR-TB treatment duration to ≤ 5 months with no relapse in the murine TB model. The Bangladesh regimen is not commonly used in some places partly due to high prevalence of resistance to FQs and PZA in DR-TB patients (17,24), especially in countries with high TB burden (25,26), for example, China, and their roles in the regimen have not been fully elucidated in an animal model. Therefore, considering the severity of resistance to FQs (16) and PZA (17–19) and the serious side effects and inconvenience associated with injectable drugs, we evaluated the effect of partially or totally replacing them in the drug combinations with the repurposed LZD to study their benefits in the new regimens based on LZD + CLF + TB47 as a backbone.

Rather than holding cohorts of mice for six months prior to relapse assessment as described before (11), here, mice were held for 3.5 months followed by two weekly immunosuppression treatment with DXM and the relapse results were comparable to our previous study (11). We propose that four months, including immunosuppression after treatment cessation, are long enough to monitor relapse, though the half-life of CLF is long and may interfere with relapse evaluation.

Here, we incorporated TB47 + CLF into all test regimens to further study regimens containing this suite of compounds. The regimen containing NCT showed very good bactericidal and sterilizing activities (Table 2 and Table 3), which may cure TB in 6 months and could possibly form an all-oral pan-TB regimen for any form of DR-TB. Inclusion of PZA in a combination regimen has been thought to even further shorten the course of TB treatment provided the infecting strain remains PZA-susceptible (27). Based on NCT, adding PZA could sharply shorten the duration further by two months or more (Table 3). This points to the remarkable role of PZA in combination with NCT, and PZA should be used when necessary.

As for LVX, it exhibited both bactericidal and sterilizing benefits in the regimen 2ALEZCT/3LEZCT over 2AEZCT/4EZCT (Figure 1, Table 2, Table 3). However, both the bactericidal and sterilizing activities of NCT/3CT and 3LNCT/3LCT, which contained NCT were indifferent (*P* > 0.05), even though they showed 6.67% and 0% relapse rates, respectively after 5 months of treatment. Adding LVX into NCT-containing regimens used here had only a limited or even no additive effect. Our data supports that notion that LVX and other FQs may be removable in such regimens. However, if LVX should be withdrawn from the Bangladesh regimen containing TB47 (ALEZCT), administration of the regimen would be subject to an extension by one month.

Because CLF + LZD + TB47 were prepared and administered together in this experiment, we speculate that in all LZD-containing regimens, had the administration of LZD been separated from administration of its companion drugs by at least 4 hours (27), their bactericidal effects might have been better than what was observed here as LZD affects the absorption of companion agents (28).

It is noticeable that “BPaL” has shown remarkably promising effect against both MDR- and XDR-TB in a phase III clinical trial (NCT02333799) by curing > 90% DR-TB patients in 6 months, though most patients have had a reduction in dose or interruption of LZD during treatment. However, side effects of the drug have been shown to be dependent on dose and duration of treatment and were manageable by withdrawal of LZD or interruption and resumption of treatment without sabotaging the efficacy of the regimen (20). Those justify the withdrawal of LZD from LZD-containing regimens after the first three months of treatment in this study. Besides the side effects, cost and accessibility of the “BPaL” regimen is also a cause for concern by many TB patients.

In conclusion, we report potentially effective all-oral regimens based on TB47 + LZD + CLF, which could achieve complete sterilization after 6 months of treatment and even 4 months with addition of PZA in a mouse TB model. The benefit of adding LVX was not quite obvious, especially in the NCT-based regimens, likely because its activity was masked by other companion agents. Though it is too early to infer that the early bactericidal and sterilizing potentials of the regimens in this study will necessarily translate directly to human TB, the promising potentials of the regimens warrant further studies to ascertain their clinical applicability in treatment of MDR- or even XDR-TB.

## MATERIALS AND METHODS

### Ethical approval

All animal procedures were approved by the Institutional Animal Care and Use Committee of the Guangzhou Institutes of Biomedicine and Health, Chinese Academy of Sciences (#2018053).

### Mycobacterial strains and growth conditions

A mouse-passaged *M. tuberculosis* H37Rv strain (ATCC 27294) preserved at −80°C was subcultured in Middlebrook 7H9 broth (Difco, Detroit, MI, USA) supplemented with 10% oleic acid-albumin-dextrose-catalase enrichment medium (BBL, Sparks, MD, USA), 0.2% glycerol, and 0.05% Tween 80 at 37°C. Broth cultures of *M. tuberculosis* at the logarithmic phase with OD_600_ between 0.8 and 1.0 were used for the infection.

### Antimicrobials

All drugs used in the study were purchased from Meilun bio (Dalian, China) except for TB47 and DXM. TB47 was synthesized and supplied by Shanghai Abotchem Co. Ltd (Shanghai, China) with a purity of ≥ 95%. DXM was purchased from Sigma-Aldrich (St. Louis, MO, USA). AMK and DXM were dissolved in sterile phosphate-buffered saline (PBS). LVX, EMB, and PZA were dissolved in distilled water. LZD, CLF, and TB47 were suspended in 0.4% sodium carboxymethyl cellulose (Solarbio, Beijing, China). All drug solutions/suspensions were prepared to deliver the drug(s) and dose(s) in a total volume of 0.2 mL per gavage or 0.1 mL per injection, based on the average mouse body weight of 20 g. Drug solutions and suspensions were prepared weekly and stored at 4°C prior to use.

### Aerosol infection and determination of bacterial burden

Five to six weeks old female BALB/c mice were purchased from Charles River Laboratories (Beijing, China). Following several days of acclimatization, 362 mice were infected with *M. tuberculosis* H37Rv at a high dose by an inhalation exposure system (Glas-Col, Terre Haute, IN, USA). The mice were block randomized into treatment groups and timed sacrifice cohorts before the start of treatment. Nine mice with 3 mice from each infection run were humanely sacrificed 1 day after infection (Day -15) and on the day of treatment initiation (Day 0) to determine the CFU counts of bacteria implanted in the lungs and at the start of treatment, respectively. *In vivo* bactericidal activities of regimens were measured by lung CFU counts during treatment for each time point.

### Chemotherapy

Drugs were prepared as previously described (11,27). The daily doses (mg/kg) are as follows: AMK 100, LVX 300, EMB 100, PZA 150, CLF 25, LZD 100 and TB47 50. The scheme is shown in Table 1. Fifteen days after infection, mice were treated once daily, 5 days per week (from Monday to Friday). AMK was administered subcutaneously, and other drugs were administered orally by gavage. LVX + EMB or CLF + LZD + TB47 were prepared and administered together in a single dose. PZA was prepared and administered alone. The lung CFU count for each mouse was determined as scheduled. Mice used for relapse assessment were subcutaneously injected with DXM (10 mg/kg) 3.5 months post-treatment cessation for immunosuppression. DXM was administered again a week later, after which the mice were kept for a week prior to sacrifice.

### Assessment of treatment efficacy

Efficacy was assessed on the basis of lung CFU counts at selected time points during treatment (a measure of bactericidal activity) and the proportion of mice with culture-positive relapse 4 months after treatment completion (a measure of sterilizing activity). Lungs were dissected from the mice and homogenized in 2.0 mL sterile PBS using glass tissue grinders. Undiluted as well as 10-fold serial dilutions of the homogenates were prepared and cultured with a volume of 0.5 mL per 7H11 agar plate. Since CLF carryover can affect the growth of *M. tuberculosis*, lung homogenates from CLF-containing groups were plated on 7H11 agar containing 0.4% (weight/volume) activated charcoal to adsorb residual CLF (29). Plates were incubated for 4 to 5 weeks at 37°C before CFU counting. Relapses were assessed 4 months (including two immunosuppressions) post-treatment cessation by equally plating the homogenate of the whole lung on 4 plates. If ≥ 1 colony appeared, the mouse was considered as relapse.

### Statistical analysis

CFU counts (x) were log-transformed to log_10_ (x + 1) prior statistical analyses, using 0.05 significance level and 95 % confidence interval.

Bactericidal activities of all regimens were compared by two-way analysis of variance with Tukey’s post-test to correct for multiple comparisons. Sterilizing activities of all the regimens were analyzed by Fisher’s exact test, adjusting for multiple comparisons. GraphPad Prism version 8.0.2 (GrahPad, San Diego, CA) was used for all statistical analyses.

## ACKNOWLEDGMENTS

This work was supported by the National Mega-project of China for Innovative Drugs (2019ZX09721001-003-003), by the National Natural Science Foundation of China (NSFC,81973372, 21920102003), by Joint Research of the Russian Science Foundation-NSFC Collaboration (21-45-00018 & 82061138019), by the Health Research Council of New Zealand-NSFC Biomedical Collaboration Fund (20/1211 & 8206112800),the Chinese Academy of Sciences Grants (154144KYSB20190005), the Grant (SKLRD-OP-201919, SKLRD-OP-202113) from the State Key Laboratory of Respiratory Disease and First Affiliated Hospital of Guangzhou Medical University. This work was sponsored by Science and Technology Innovation Leader of Guangdong Province (2016TX03R095 to TZ), the UCAS (to Y.B.) and “One Belt One Road” (to A.M.S.) Master Fellowship Programs for international students, CAS-TWAS President’s PhD Fellowship Program (to G.C.) for international students, and the Postdoctoral Fellowship from the University of Chinese Academy of Sciences (to H.M.A.H). The funders had no role in study design, data collection and analysis, decision to publish, or preparation of the manuscript.

